# Comparative Analysis of Emerging B.1.1.7+E484K SARS-CoV-2 isolates from Pennsylvania

**DOI:** 10.1101/2021.04.21.440801

**Authors:** Ahmed M. Moustafa, Colleen Bianco, Lidiya Denu, Azad Ahmed, Brandy Neide, John Everett, Shantan Reddy, Emilie Rabut, Jasmine Deseignora, Michael D. Feldman, Kyle G. Rodino, Frederic Bushman, Rebecca M. Harris, Josh Chang Mell, Paul J. Planet

## Abstract

Rapid whole genome sequencing of SARS-CoV-2 has presented the ability to detect new emerging variants of concern in near real time. Here we report the genome of a virus isolated in Pennsylvania in March 2021 that was identified as lineage B.1.1.7 (VOC-202012/01) that also harbors the E484K spike mutation, which has been shown to promote “escape” from neutralizing antibodies *in vitro*. We compare this sequence to the only 5 other B.1.1.7+E484K genomes from Pennsylvania, all of which were isolated in mid March. Beginning in February 2021, only a small number (n=60) of isolates with this profile have been detected in the US, and only a total of 253 have been reported globally (first in the UK in December 2020). Comparative genomics of all currently available high coverage B.1.1.7+E484K genomes (n=235) available on GISAID suggested the existence of 7 distinct groups or clonal complexes (CC; as defined by GNUVID) bearing the E484K mutation raising the possibility of 7 independent acquisitions of the E484K spike mutation in each background. Phylogenetic analysis suggested the presence of at least 3 distinct clades of B.1.1.7+E484K circulating in the US, with the Pennsylvanian isolates belonging to two distinct clades. Increased genomic surveillance will be crucial for detection of emerging variants of concern that can escape natural and vaccine induced immunity.

---

During the past six months of the pandemic several variants of concern (VOC), each represented by a constellation of specific mutations thought to enhance viral fitness, have emerged in viral lineages from the UK (20I/501Y.V1; B.1.1.7), South Africa (20H/501Y.V2; B.1.351), and Brazil (20J/501Y.V3; P.1). These lineages were concerning due to likely increased transmission rates^1–6^. Two of these lineages, B.1.351 and P.1 were of specific concern because they harbor the mutation E484K, which has been shown to enhance escape from neutralizing antibody inhibition in vitro^7^, and may be associated with reduced efficacy of the vaccine^8–11^. In general, viruses from the B.1.1.7 lineage do not harbor this mutation. However, in February 2021 Public Health England (PHE) published a concerning report of eleven B.1.1.7 genomes that had acquired the E484K spike mutation^12^.

Here we report a B.1.1.7 isolate with the E484K spike mutation isolated in southeastern Pennsylvania (PA). Our laboratory at the Children’s hospital of Philadelphia performed sequencing on randomly selected isolates collected since January 2021. **Figure 1A** shows the diversity of 114 randomly sequenced genomes. Lineages B.1.1.7, B.1.429 (California), B.1.526 (New York) and R.1 (International lineage with the E484K mutation) accounted for 69% of the sequenced genomes in March. There was a massive increase in lineage B.1.1.7 from 2% (1/47) in February to 42% in March (15/36). Interestingly, one B.1.1.7 isolate carried the E484K spike mutation that is present in the South African and Brazilian lineages.

**Figure 1.**
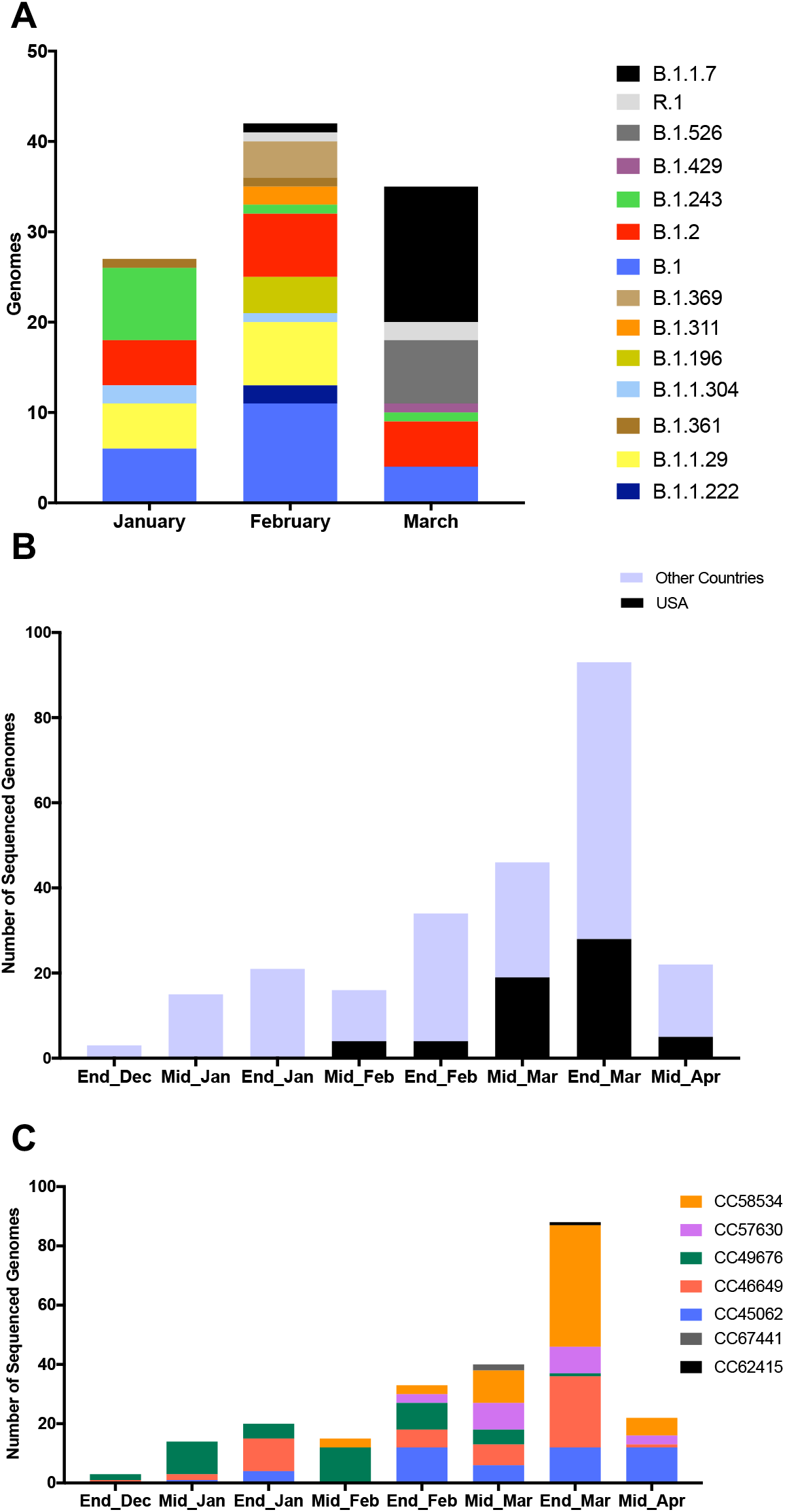
Diversity of SARS-CoV-2 in Philadelphia and global diversity of sequenced B.1.1.7+E484K genomes. **A.** Stacked bar plot showing the diversity of random genomes sequenced by our laboratory at Children’s Hospital of Philadelphia during January, February and March 2021. Ten lineages that were represented by only one genome (B.1.1, B.1.1.106, B.1.1.129, B.1.1.197, B.1.1.281, B.1.1.296, B.1.119, B.1.234, B.1.350, B.1.409) were excluded from the plot. One isolate that is B.1.526.1 was counted with the parent B.1.526 for easier visualization. **B.** Bar plot showing number of GISAID genomes (n=250) that are 20I/501Y.V1 and have the E484K spike mutation over time in the US and globally. **C.** Diversity of 236 isolates according to GNUVID. Bar plot showing relative abundance of circulating clonal complexes (CC) for the 236 B.1.1.7+E484K isolates (typed by GNUVID). The bar plot shows that the isolates belong to 7 different CCs. Isolate EPI_ISL_1385215 was not assigned to any of the 7 CCs (CC255). Fourteen isolates were excluded from the plot as they had > 5% nucleotides designated “N” in the sequence.

To better understand the relationship between this isolate and publicly available SARS-CoV-2 genomes, we compared it to all available B.1.1.7+E484K high coverage genomes available on GISAID^13^ (n=235). Since the first report by PHE in February, a total of 253 B.1.1.7+E484K genomes have been uploaded to GISAID from England and 14 other countries (Germany, France, Italy, Poland, Sweden, Ireland, Netherlands, Portugal, Wales, Turkey, Slovakia, Austria, Czech Republic and USA)^13^ (as of 04/17/2021).

A temporal plot of the number of B.1.1.7+E484K isolates collected between December 2020 to March 2021 (2-week window) is shown in **Figure 1B**. The first isolate of the 60 US isolates available on GISAID was collected on 02/06/2021 from Oregon (OR). Isolates were also reported from 15 other states (New York, North Carolina, Connecticut, Georgia, New Jersey, Maryland, Florida, West Virginia, California, Pennsylvania, Michigan, Texas, Massachusetts, Washington, and Colorado). Of these isolates 48% were from Florida (n=17) and New York (n=12) and 28% were from New Jersey (n=7), California (n=4) and Pennsylvania (n=6). Two isolates were from Oregon (OR), Connecticut (CT), Maryland (MD), and single isolates are recorded from Georgia (GA), Texas (TX), Massachusetts (MA), Washington (WA), Colorado (CO), West Virginia (WV), Michigan (MI), and North Carolina (NC). The number of US isolates in March (n=47 including the PA isolates) was nearly 6 times the number of the isolates reported in February. This increase raises the concern that more B.1.1.7+E484K sequences may be emerging even as herd immunity increases by natural immunity and vaccines.

Although all 236 genomes were typed as B.1.1.7 using Pangolin^14^, a more granular view using our typing tool “GNUVID”^15^ shows that they belong to 7 different clonal complexes (CCs 45062, 46649, 49676, 57630, 58534, 62415 and 67441) (**Figure 1C and Supplementary Table 1**). In the GNUVID typing system, these correspond to 7 of 10 CCs in the B.1.1.7 lineage. For each of these CCs, representative sequences without the E484K mutation have been circulating since at least November 2020, predating the first E484K in each CC. This raises the possibility that the E484K mutation was acquired independently in each of these CCs in independent events.

Phylogenetic analysis of the 235 B.1.1.7+E484K GISAID isolates showed that US isolates are found in at least 3 different clades. The genome presented here falls in a well-supported clade of 28 isolates, 6 of which were from the US (CT, FL, OR, PA and NY), 18 from Sweden, 2 from Poland and 1 from Germany (**Figure 2A**). The only other 4 isolates reported from PA, were in a large clade containing the majority of US genomes, and were located in a well-supported subclade with genomes from the nearby state of West Virginia.

**Figure 2.**
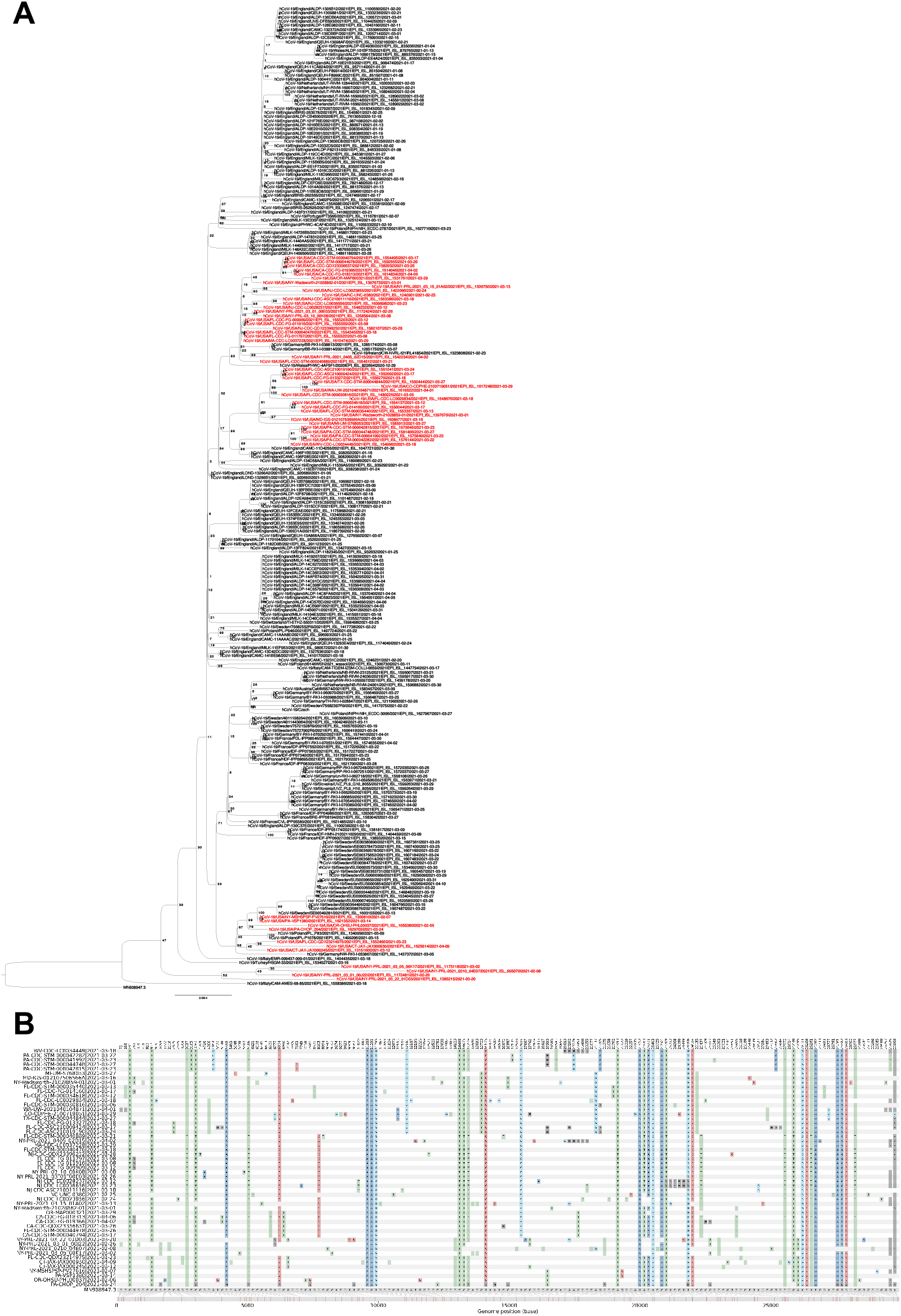
SNP-based Phylogeny and variations of the B.1.1.7+E484K isolates. **A.** Maximum likelihood tree of the B.1.1.7+E484K isolates. US isolates are in red. For the CHOP_204 isolate the alternative allele was called as consensus if its frequency was at least 0.75. The tree was rooted with MN908947.3. Bootstrap values are shown on the branches. **B.** SNP patterns in the 53 US isolates compared to MN908947.3. SNP variations in the 236 isolates are shown in Supplementary Figure 1. Mutations identified in CHOP_204 are available in Supplementary Table 2. Seven US isolates were excluded from the plot as they had > 5% nucleotides designated “N” in the sequence. An acknowledgement table of the submitting laboratories providing the SARS-CoV-2 genomes used in this study is in Supplemental Table 3.

Analysis of SNPs in the 236 isolates compared to the reference MN908947.3^16^ (**Figure 2B and Supplementary Figure 1**) showed that the isolate presented here had 12/17 of the B.1.1.7 defining SNPs (**Supplementary Table 2**), while the other Pennsylvanian isolate in the same clade had 17/17 of the SNPs. It also shared with 9 other US isolates a stop mutation (A28095T) in ORF8 (**Figure 2B**).

Here we present a comparative analysis of the first SARS-CoV-2 B.1.1.7 isolates detected in PA that harbor the E484K spike mutation, a mutation that could be associated with reduced efficacy of both vaccine-induced and natural immunity. Our analysis suggests that multiple lineages of B.1.1.7+E484K are circulating in the US, and that these lineages may have acquired E484K independently.

## Methods

A nasopharyngeal swab sample that had residual volume after initial laboratory processing, positive PCR testing for SARS-CoV-2, was obtained for this study. RNA was extracted from nasopharyngeal swab samples using QIAamp Viral RNA Mini (Qiagen). Whole genome sequencing was done by The Genomics Core Facility at Drexel University. Briefly, WGS of extracted viral RNA was performed as previously described using Paragon Genomics CleanPlex SARS-CoV-2 Research and Surveillance NGS Panel^17,18^. Libraries were quantified using the Qubit dsDNA HS (High Sensitivity) Assay Kit (Invitrogen) with the Qubit Fluorometer (Invitrogen). Library quality was assessed using Agilent High Sensitivity DNA Kit and the 2100 Bioanalyzer instrument (Agilent). Libraries were then normalized to 5nM and pooled in equimolar concentrations. The resulting pool was quantified again using the Qubit dsDNA HS (High Sensitivity) Assay Kit (Invitrogen) and diluted to a final concentration of 4nM; libraries were denatured and diluted according to Illumina protocols and loaded on the MiSeq at 10pM. Paired-end and dual-indexed 2×150bp sequencing was done using MiSeq Reagent Kits v3 (300 cycles). Sequences were demultiplexed and basecalls were converted to FASTQ using bcl2fastq2 v2.20. The FASTQ reads were then processed to consensus sequence and variants were identified using the ncov2019-artic-nf pipeline (https://github.com/connor-lab/ncov2019-artic-nf). Briefly, the pipeline uses iVar^19^ for primer trimming and consensus sequence making (options: -- ivarFreqThreshold 0.75). A bed file for the Paragon kit primers was used in the pipeline.

All 253 SARS-CoV-2 genomes that were assigned to Pango lineage^14^ B.1.1.7 and possessing the E484K spike mutation (including the study isolate CHOP_204) were downloaded from GISAID^13^ on 04/17/2021. An acknowledgement table of the submitting laboratories providing the SARS-CoV-2 genomes used in this study is in **Supplemental Table 3**. Seventeen sequences were excluded for lower coverage (> 5% Ns) (n=14) and missing collection date (n=3). All the high coverage SARS-CoV-2 genomes (n=236) were assigned a clonal complex using the GNUVID v2.2 database (version January 6^th^ 2021)^15^. Temporal plots were plotted in GraphPad Prism v7.0a.

To show the relationship amongst the genomes of the 236 isolates, a maximum likelihood tree was constructed. Briefly, consensus SARS-CoV-2 sequences for the 236 isolates were aligned to MN908947.3^16^ using MAFFT’s FFT-NS-2 algorithm ^20^ (options: --add --keeplength)). The 5’ and 3’ untranslated regions were masked in the alignment file using a custom script. A maximum likelihood tree using IQ-TREE 2^21^ was then estimated using the GTR+F+I model of nucleotide substitution^22^, default heuristic search options, and ultrafast bootstrapping with 1000 replicates^23^. The tree was rooted to MN908947.3. The snipit tool was then used to summarize the SNPs in the 236 isolates relative to MN908947.3 (https://github.com/aineniamh/snipit).

The sample was obtained by as part of routine clinical care, solely for non-research purposes, carrying minimal risk, and were therefore granted a waiver of informed consent as reviewed under protocol number under IRB 21-018478.

## Availability of data and material

The sequence has been uploaded to GISAID with accession number EPI_ISL_1629709.

## Acknowledgements

We would like to thank the Global Initiative on Sharing All Influenza Data (GISAID) and thousands of contributing laboratories for making the genomes publicly available. A full acknowledgements table is available in Supplementary Table 3. We would like to acknowledge the staff members of the Drexel Genomics Core Facility at the Drexel University College of Medicine for processing and sequencing the isolates. P.J.P and A.M.M are supported by 1R01AI137526-01 and 1R21AI144561-01A1 (A.M.M. and P.J.P.), and R01NR015639 (P.J.P.).

---

**Supplementary Table 1. Excel Sheet of GNUVID results for the 236 isolates.**

**Supplementary Table 2.** Mutations and deletions in CHOP_204 compared to MN908947.3.

| **Mutation** | **Protein** | **AA change** | **Frequency** |
| --- | --- | --- | --- |
| C241T | - | - | 1 |
| C913T | ORF1ab | synonymous | 0.92 |
| C1059T | ORF1ab | T265I | 0.33 |
| C2110T | ORF1ab | synonymous | 0.98 |
| C3037T | ORF1ab | synonymous | 1 |
| C3267T | ORF1ab | T1001I | 0.64 |
| C4320T | ORF1ab | synonymous | 0.38 |
| C5388A | ORF1ab | A1708D | 0.65 |
| C5986T | ORF1ab | synonymous | 0.62 |
| T6954C | ORF1ab | I2230T | 0.74 |
| T7984C | ORF1ab | synonymous | 0.65 |
| T9867C | ORF1ab | L3201P | 0.33 |
| 11288 (del-9) | ORF1ab | SGF3675-77 deletion | 0.99 |
| C12781T | ORF1ab | synonymous | 0.96 |
| C14120T | ORF1ab | Q4619* | 0.95 |
| C14408T | ORF1ab | synonymous | 1 |
| C14676T | ORF1ab | P4804L | 0.96 |
| C15279T | ORF1ab | T5005I | 0.66 |
| T16176C | ORF1ab | L5304P | 0.72 |
| A16500C | ORF1ab | K5412T | 0.30 |
| C16887T | ORF1ab | synonymous | 0.31 |
| C19390T | ORF1ab | synonymous | 1 |
| C21575T | S | L5F | 0.35 |
| 21765 (del6) | S | HV69-70 deletion | 0.99 |
| 21991 (del3) | S | Y144 deletion | 0.98 |
| G23012A | S | E484K | 0.77 |
| A23063T | S | N501Y | 0.95 |
| C23271A | S | A570D | 1 |
| A23403G | S | D614G | 1 |
| C23604A | S | P681H | 0.98 |
| C23664T | S | A701V | 0.41 |
| C23709T | S | T716I | 0.99 |
| T24506G | S | S982A | 0.55 |
| G24914C | S | D1118H | 0.94 |
| C25517T | ORF3a | P42L | 0.36 |
| C27972T | ORF8 | Q27* | 0.93 |
| A28095T | ORF8 | K68* | 0.93 |
| A28111G | ORF8 | Y73C | 0.96 |
| A28271 (del1) | - | deletion | 0.97 |
| GAT28280CTA | N | D3L | 0.97 |
| C28869T | N | P199L | 0.38 |
| GGG28881AAC | N | R203K, G204R | 0.55 |
| C28977T | N | S235F | 0.88 |
| C29137T | N | synonymous | 0.54 |

**Supplementary Table 3. GISAID Acknowledgement Table.**

**Supplementary Figure 1. SNP variations in all available 20I/501Y.V1+E484K isolates.**

