## Supplementary figures and images for "Comparative Analysis of Emerging B.1.1.7+E484K SARS-CoV-2 isolates from Pennsylvania"

### Supplementary Figure 1

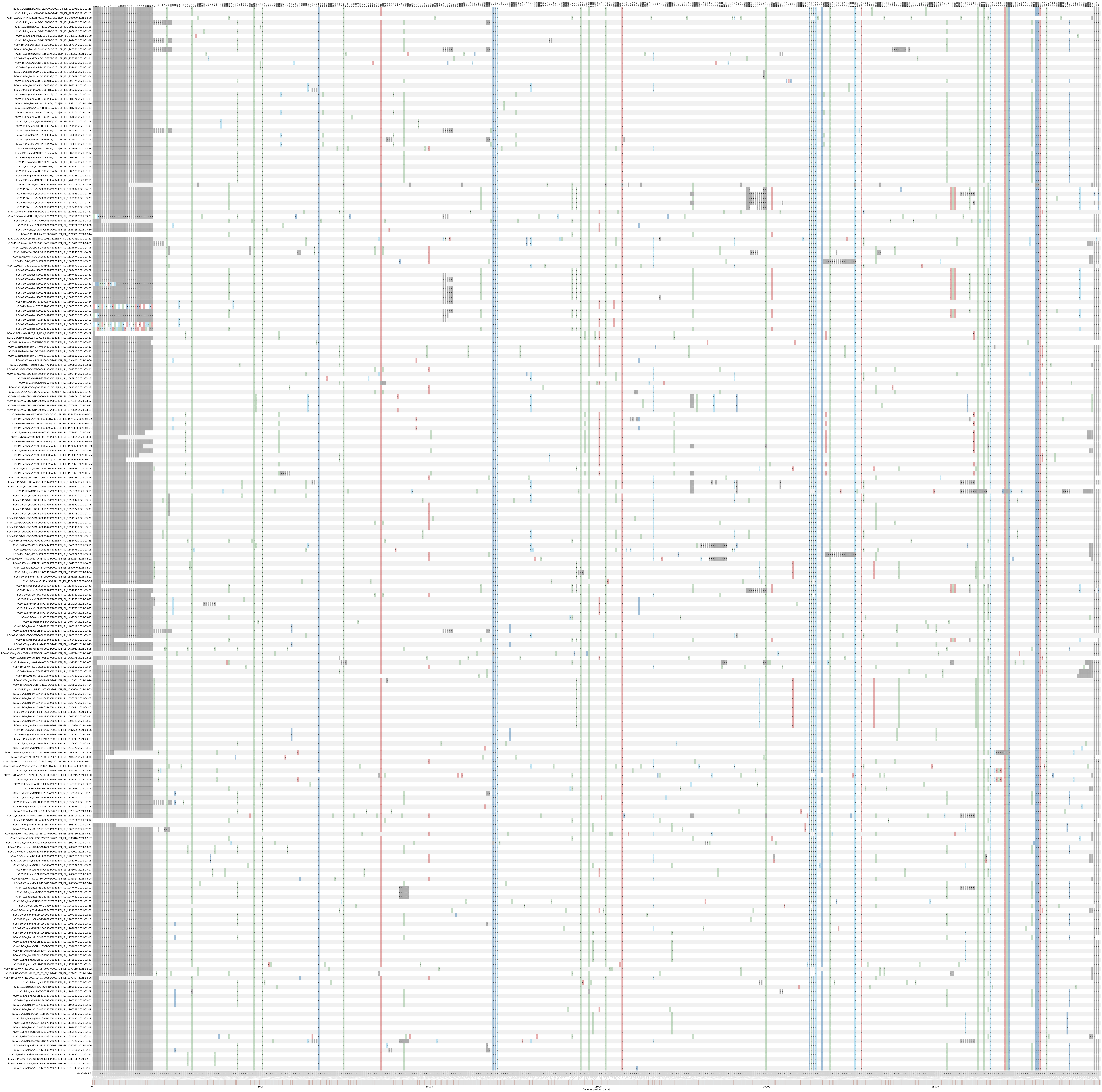
