## Supplementary Table 3 for "Comparative Analysis of Emerging B.1.1.7+E484K SARS-CoV-2 isolates from Pennsylvania"

We gratefully acknowledge the following Authors from the Originating laboratories responsible for obtaining the specimens, as well as the Submitting laboratories where the genome data were generated and shared via GISAID, on which this research is based.

All Submitters of data may be contacted directly via www.gisaid.org

Authors are sorted alphabetically.

| Accession ID | Originating Laboratory | Submitting Laboratory | Authors |
|---|---|---|---|
| EPI_ISL_1018343 | Lighthouse Lab in Alderley Park | Wellcome Sanger Institute for the COVID-19 Genomics UK (COG-UK) Consortium | Jacquelyn Wynn, Mairead Hyland, The Lighthouse Lab in Alderley Park and Alex Alderton, Roberto Amato, Jeffrey Barrett, Sonia Goncalves, Ewan Harrison, David K. Jackson, Ian Johnston, Dominic Kwiatkowski, Cordelia Langford, John Sillitoe on behalf of the Wellcome Sanger Institute COVID-19 Surveillance Team |
| EPI_ISL_1020302 | Dutch COVID-19 response team | National Institute for Public Health and the Environment (RIVM) | Adam Meijer, Harry Vennema, Dirk Eggink, Jeroen Cremer, Sharon van den Brink, Bas van der Veer, AnneMarie van den Brandt, Florian Zwagemaker, Dennis Schmitz, Chantal Reusken, on behalf of the national COVID-19 response team |
| EPI_ISL_1045160 | Lighthouse Lab in Alderley Park | Wellcome Sanger Institute for the COVID-19 Genomics UK (COG-UK) Consortium | Jacquelyn Wynn, Mairead Hyland, The Lighthouse Lab in Alderley Park and Alex Alderton, Roberto Amato, Jeffrey Barrett, Sonia Goncalves, Ewan Harrison, David K. Jackson, Ian Johnston, Dominic Kwiatkowski, Cordelia Langford, John Sillitoe on behalf of the Wellcome Sanger Institute COVID-19 Surveillance Team |
| EPI_ISL_1045593 | Lighthouse Lab in Milton Keynes | Wellcome Sanger Institute for the COVID-19 Genomics UK (COG-UK) Consortium | The Lighthouse Lab in Milton Keynes and Alex Alderton, Roberto Amato, Jeffrey Barrett, Sonia Goncalves, Ewan Harrison, David K. Jackson, Ian Johnston, Dominic Kwiatkowski, Cordelia Langford, John Sillitoe on behalf of the Wellcome Sanger Institute COVID-19 Surveillance Team |
| EPI_ISL_1047721 | Virology Department, Sheffield Teaching Hospitals NHS Foundation Trust/Department of Infection, Immunity and Cardiovascular Disease, The Medical School, University of Sheffield | COVID-19 Genomics UK (COG-UK) Consortium | Thushan de Silva, Matthew Parker, Nikki Smith, Adri Angyal, Rebecca Brown, Luke Green, Rachel Tucker, Paul Parsons, Danielle Groves, Katie Johnson, Laura Carrilero, Alex Keeley, Dave Partridge, Matthew Wyles, Benjamin Lindsey, Mehmet Yavuz, Mohammad Raza, Cariad Evans |
| EPI_ISL_1055380 | Oregon State Public Health Laboratory | Oregon SARS-CoV-2 Genome Sequencing Center | Brendan L. O'Connell, Sally Grindstaff, Kayla Carter, Sonia Acharya, John Fontana, LaDonna Grenz, Ruth V. Nichols, Alec J. Hirsch, Donna Hansel, Guang Fan, Xuan Qin, Daniel N. Streblow, William B. Messer, Andrew C. Adey, Benjamin N. Bimber, Brian J. O'Roak |
| EPI_ISL_1069921 | Lighthouse Lab in Glasgow | Wellcome Sanger Institute for the COVID-19 Genomics UK (COG-UK) Consortium | Harper VanSteenhouse, Yumi Kasai, David Gray, Carol Clugston, Anna Dominiczak and Alex Alderton, Roberto Amato, Jeffrey Barrett, Sonia Goncalves, Ewan Harrison, David K. Jackson, Ian Johnston, Dominic Kwiatkowski, Cordelia Langford, John Sillitoe on behalf of the Wellcome Sanger Institute COVID-19 Surveillance Team |
| EPI_ISL_1089490 | Dutch COVID-19 response team | National Institute for Public Health and the Environment (RIVM) | Adam Meijer, Harry Vennema, Dirk Eggink, Jeroen Cremer, Sharon van den Brink, Bas van der Veer, AnneMarie van den Brandt, Florian Zwagemaker, Dennis Schmitz, Chantal Reusken, on behalf of the national COVID-19 response team |
| EPI_ISL_1100238, EPI_ISL_1100560, EPI_ISL_1101487 | Lighthouse Lab in Alderley Park | Wellcome Sanger Institute for the COVID-19 Genomics UK (COG-UK) Consortium | Jacquelyn Wynn, Mairead Hyland, The Lighthouse Lab in Alderley Park and Alex Alderton, Roberto Amato, Jeffrey Barrett, Sonia Goncalves, Ewan Harrison, David K. Jackson, Ian Johnston, Dominic Kwiatkowski, Cordelia Langford, John Sillitoe on behalf of the Wellcome Sanger Institute COVID-19 Surveillance Team |
| EPI_ISL_1104425 | Liverpool Clinical Laboratories | COVID-19 Genomics UK (COG-UK) Consortium | Sam Haldenby, Anita Lucaci, Steve Paterson, Julian Hiscox, Alistair Darby, M Almsaud, A Alrezaihi, Muhannad Alruwaili, Stuart D Armstrong, Jones Benjamin, Eleanor G Bentley, Anu Chawla, Jordan J Clark, Angela Cowell, Richard Eccles, Isabel García-Dorival, Matthew Gemmell, Alessandro Gerada, PKF Gilmore, Richard Gregory, Ximeng Han, Catherine Hartley, Margaret Hughes, Miren Iturriza-Gomara, James Johnson, L Luu, Jenifer Manson, Charlotte Nelson, Elaine O'Toole, Cassie Olateju, Rebekah Penrice-Randal , Lucille Rainbow, N.P Randle, Trevor Ian Robinson, Parul Sharma, Ghada T Shawli, James P Stewart, Neil Swainston, Ecaterina Vamos, Joanne Watts, Mark Whitehead |
| EPI_ISL_1105933 | Wales Specialist Virology Centre Sequencing lab: Pathogen Genomics Unit | Public Health Wales Microbiology Cardiff Wales Specialist Virology Centre | Catherine Moore, Johnathan Evans, Laura Gifford, Malorie Perry, Simon Cottrell, Angela Marchbank, Alec Birchley, Alexander Adams, Amy Gaskin, Bree Gatica-Wilcox, Jason Coombes, Joel Southgate, Lauren Gilbert, Lee Graham, Nicole Pacchiarini, Sara Kumziene-Summerhayes, Sarah Taylor, Sophie Jones, Sara Rey, Matthew Bull, Joanne Watkins, Sally Corden, Tom Connor |
| EPI_ISL_1114929 | Lighthouse Lab in Alderley Park | Wellcome Sanger Institute for the COVID-19 Genomics UK (COG-UK) Consortium | Jacquelyn Wynn, Mairead Hyland, The Lighthouse Lab in Alderley Park and Alex Alderton, Roberto Amato, Jeffrey Barrett, Sonia Goncalves, Ewan Harrison, David K. Jackson, Ian Johnston, Dominic Kwiatkowski, Cordelia Langford, John Sillitoe on behalf of the Wellcome Sanger Institute COVID-19 Surveillance Team |
| EPI_ISL_1116781 | Instituto Nacional de Saude (INSA) | Instituto Nacional de Saude (INSA) | Borges et al |
| EPI_ISL_1172424, EPI_ISL_1172481, EPI_ISL_1173118 | Pandemic Response Lab - NYC | Pandemic Response Lab, R&D | Henry Lee, Michael Hammerling, Melissa Hopkins, Cybill del Castillo, Shinyoung Clair Kang, William Ward, Pradeep Bugga, Haiping Hao, Jon Laurent |
| EPI_ISL_1174049, EPI_ISL_1175868 | Lighthouse Lab in Glasgow | Wellcome Sanger Institute for the COVID-19 Genomics UK (COG-UK) Consortium | Harper VanSteenhouse, Yumi Kasai, David Gray, Carol Clugston, Anna Dominiczak and Alex Alderton, Roberto Amato, Jeffrey Barrett, Sonia Goncalves, Ewan Harrison, David K. Jackson, Ian Johnston, Dominic Kwiatkowski, Cordelia Langford, John Sillitoe on behalf of the Wellcome Sanger Institute COVID-19 Surveillance Team |
| EPI_ISL_1176993 | Northumbria University / South Tees Hospitals NHS Foundation Trust / North Cumbria Integrated Care NHS Foundation Trust / North Tees and Hartlepool NHS Foundation Trust / Newcastle Hospitals NHS Foundation Trust | COVID-19 Genomics UK (COG-UK) Consortium | Darren L Smith,Andrew Nelson,Matthew Bashton,Greg R Young,Joshua Loh,John Allan,Mohammad A Tariq,Giles S Holt,Gary Black,Wen C Yew,Lynn Dover,Paul Baker,Steve Liggett,Sarah Essex,Jane Greenaway,Debra Padgett,Clive Graham,Garren Scott,Edward Barton,Emma Swindells,Brendan Payne,Jennifer Collins,Yusri Taha,Gary Eltringham |
| EPI_ISL_1186598, EPI_ISL_1186739, EPI_ISL_1189089, EPI_ISL_1205714, EPI_ISL_1205721 | Lighthouse Lab in Alderley Park | Wellcome Sanger Institute for the COVID-19 Genomics UK (COG-UK) Consortium | Jacquelyn Wynn, Mairead Hyland, The Lighthouse Lab in Alderley Park and Alex Alderton, Roberto Amato, Jeffrey Barrett, Sonia Goncalves, Ewan Harrison, David K. Jackson, Ian Johnston, Dominic Kwiatkowski, Cordelia Langford, John Sillitoe on behalf of the Wellcome Sanger Institute COVID-19 Surveillance Team |
| EPI_ISL_1206501 | Lighthouse Lab in Cambridge | Wellcome Sanger Institute for the COVID-19 Genomics UK (COG-UK) Consortium | Rob Howes, The Lighthouse Lab in Cambridge and Alex Alderton, Roberto Amato, Jeffrey Barrett, Sonia Goncalves, Ewan Harrison, David K. Jackson, Ian Johnston, Dominic Kwiatkowski, Cordelia Langford, John Sillitoe on behalf of the Wellcome Sanger Institute COVID-19 Surveillance Team |
| EPI_ISL_1207256 | Lighthouse Lab in Alderley Park | Wellcome Sanger Institute for the COVID-19 Genomics UK (COG-UK) Consortium | Jacquelyn Wynn, Mairead Hyland, The Lighthouse Lab in Alderley Park and Alex Alderton, Roberto Amato, Jeffrey Barrett, Sonia Goncalves, Ewan Harrison, David K. Jackson, Ian Johnston, Dominic Kwiatkowski, Cordelia Langford, John Sillitoe on behalf of the Wellcome Sanger Institute COVID-19 Surveillance Team |
| EPI_ISL_1211960 | Dianovis GmbH Greiz | Robert Koch Institute | unknown |
| EPI_ISL_1232682 | Dutch COVID-19 response team | National Institute for Public Health and the Environment (RIVM) | Adam Meijer, Harry Vennema, Dirk Eggink, Jeroen Cremer, Sharon van den Brink, Bas van der Veer, AnneMarie van den Brandt, Florian Zwagemaker, Dennis Schmitz, Chantal Reusken, on behalf of the national COVID-19 response team |
| EPI_ISL_1240901 | Clinical Molecular Microbiology Laboratory, UNC Hospitals | Jeremy Wang | Jeremy Wang, Alexander Rubinsteyn, Colleen Rice, Jason Smedberg, Shawn Hawken, Melissa Miller, Corbin Jones, Robert Hagan |
| EPI_ISL_1245353 | Lighthouse Lab in Glasgow | Wellcome Sanger Institute for the COVID-19 Genomics UK (COG-UK) Consortium | Harper VanSteenhouse, Yumi Kasai, David Gray, Carol Clugston, Anna Dominiczak and Alex Alderton, Roberto Amato, Jeffrey Barrett, Sonia Goncalves, Ewan Harrison, David K. Jackson, Ian Johnston, Dominic Kwiatkowski, Cordelia Langford, John Sillitoe on behalf of the Wellcome Sanger Institute COVID-19 Surveillance Team |
| EPI_ISL_1246231 | Lighthouse Lab in Cambridge | Wellcome Sanger Institute for the COVID-19 Genomics UK (COG-UK) Consortium | Rob Howes, The Lighthouse Lab in Cambridge and Alex Alderton, Roberto Amato, Jeffrey Barrett, Sonia Goncalves, Ewan Harrison, David K. Jackson, Ian Johnston, Dominic Kwiatkowski, Cordelia Langford, John Sillitoe on behalf of the Wellcome Sanger Institute COVID-19 Surveillance Team |
| EPI_ISL_1247469, EPI_ISL_1247474 | University of Exeter | COVID-19 Genomics UK (COG-UK) Consortium | Ben Temperton,Aaron Jeffries,Michelle Michelsen,Joanna Warwick-Dugdale,Audrey Farbos,Robyn Manley,Stephen Michell,Jane Masoli |

| | | | |
|---|---|---|---|
| EPI_ISL_1248154 | University College London, Great Ormond Street Hospital for Children NHS Foundation Trust, Imperial College Healthcare NHS Trust | COVID-19 Genomics UK (COG-UK) Consortium | Sergi Castellano, Rachel Williams, Mark Kristiansen, Paola Resende Silva, Sunando Roy, Tony Brooks, Helena Tutill, Paola Niola, Patricia Dyal, Charlotte Williams, Leysa Forrest, Yasmin Panchbhaya, Jacqueline Findlay, Samuel Weeks, Julianne Brown, Kathryn Harris, Paul Randell, James Price, Alison Holmes, Judith Breuer |
| EPI_ISL_1248566 | Virology Department, Sheffield Teaching Hospitals NHS Foundation Trust/Department of Infection, Immunity and Cardiovascular Disease, The Medical School, University of Sheffield | COVID-19 Genomics UK (COG-UK) Consortium | Thushan de Silva, Matthew Parker, Nikki Smith, Adri Angyal, Rebecca Brown, Luke Green, Rachel Tucker, Paul Parsons, Danielle Groves, Katie Johnson, Laura Carrilero, Alex Keeley, Dave Partridge, Matthew Wyles, Benjamin Lindsey, Mehmet Yavuz, Mohammad Raza, Cariad Evans |
| EPI_ISL_1258584, EPI_ISL_1258688 | Pandemic Response Lab - NYC | Pandemic Response Lab, R&D | Henry Lee, Michael Hammerling, Melissa Hopkins, Cybill del Castillo, Shinyoung Clair Kang, William Ward, Pradeep Bugga, Haiping Hao, Jon Laurent |
| EPI_ISL_1263057 | Labo Analyses Med | National Reference Center for Viruses of Respiratory Infections, Institut Pasteur, Paris | Marion Barbet, Sylvie Behillil, Méline Bizard, Angela Brisebarre, Camille Capel, Etienne Simon-Lorière, Vincent Enouf, Maud Vanpeene, Sylvie van der Werf,Roquebert BéNéDicte |
| EPI_ISL_1275490, EPI_ISL_1275545, EPI_ISL_1276592 | Lighthouse Lab in Glasgow | Wellcome Sanger Institute for the COVID-19 Genomics UK (COG-UK) Consortium | Harper VanSteenhouse, Yumi Kasai, David Gray, Carol Clugston, Anna Dominiczak and Alex Alderton, Roberto Amato, Jeffrey Barrett, Sonia Goncalves, Ewan Harrison, David K. Jackson, Ian Johnston, Dominic Kwiatkowski, Cordelia Langford, John Sillitoe on behalf of the Wellcome Sanger Institute COVID-19 Surveillance Team |
| EPI_ISL_1285174, EPI_ISL_1285175 | Klinikum Ernst von Bergmann gemeinnützige GmbH - stationärer Bereich | Robert Koch Institute | unknown |
| EPI_ISL_1289022, EPI_ISL_1289029 | Dutch COVID-19 response team | National Institute for Public Health and the Environment (RIVM) | Adam Meijer, Harry Vennema, Dirk Eggink, Jeroen Cremer, Sharon van den Brink, Bas van der Veer, AnneMarie van den Brandt, Florian Zwagemaker, Dennis Schmitz, Chantal Reusken, on behalf of the national COVID-19 response team |
| EPI_ISL_1298956 | ASL Napoli 1 Centro | AMES Centro Polidiagnostico Strumentale S.r.l. | "Giovanni Savarese, Raffaella Ruggiero, Eloisa Evangelista, Antonella Di Carlo, Luisa Circelli, Luigi D'Amore, Nadia Petrillo, Monica Ianniello, Roberto Sirica, Maurizio D'Amora, Antonio Fico" |
| EPI_ISL_1300730 | Wojewodzka Stacja Sanitarno-Epidemiologiczna w Olsztynie, Laboratorium Badan Epidemiologiczno-Klinicznych | Wojewodzka Stacja Sanitarno-Epidemiologiczna w Olsztynie, Laboratorium Badan Epidemiologiczno-Klinicznych | Sylwia Krzetowska, Monika Czerminska, Ewa Liszewska, Tomasz Jakubczak, Marta Lukian, Patryk Bielecki, Aleksandra Kobiatko, Emilia Tarabasz, Paulina Rozycka, Barbara Dolinska |
| EPI_ISL_1300810 | MSHS Clinical Microbiology Laboratories | MSHS Pathogen Surveillance Program | Ana S. Gonzalez-Reiche, Hala Alshammary, Mitchell J. Sullivan, Brianne Ciferri, Ajay Obla, Angela Amoako, Mahmoud Awawda, Daniel Floda, Julia Matthews, Ashley Salimbangon, Levy Sominsky, Katherine Beach, Kayla Russo, Charles Gleason, Shelcie Fabre, Giulio Kleiner, Zenab Khan, Bremy Alburquerque, Adriana van de Guchte, Komal Srivastava, Matthew M. Hernandez, Jayeeta Dutta, Denise Jurczyszak, Nancy Francoeur, Betsaida Salom Melo, Irina Oussenko, Gintaras Deikus, Juan Soto, Shwetha Hara Sridhar, Ying-Chih Wang, Kathryn Twyman, Deena R. Altman, Robert Sebra, Adolfo García-Sastre, Marta Luksza, Gopi Patel, Sarah Schaefer, Melissa Gitman, Michael D. Nowak, Alberto Paniz-Mondolfi, Emilia Mia Sordillo, Viviana Simon, Harm van Bakel |
| EPI_ISL_1306750 | Pandemic Response Lab - NYC | Pandemic Response Lab, R&D | Henry Lee, Michael Hammerling, Melissa Hopkins, Cybill del Castillo, Shinyoung Clair Kang, William Ward, Pradeep Bugga, Sol Rey, Dylan Law, Haiping Hao, Jon Laurent |
| EPI_ISL_1308159, EPI_ISL_1308177 | Quadram Institute Bioscience | COVID-19 Genomics UK (COG-UK) Consortium | Dave J. Baker, Gemma L. Kay, Alp Aydin, Thanh Le-Viet, Steven Rudder, Ana P. Tedim, Anastasia Kolyva, Maria Diaz, Leonardo de Oliveira Martins, Nabil-Fareed Alikhan, Lizzie Meadows, Rachael Stanley, Ngozi Elumogo, Muhammed Yasir, Nicholas M. Thomson, Alexander J Trotter, Rachel Gilroy, Samuel Bloomfield, Claire Stuart, Andrew Bell, Reenesh Prakash, Samir Dervisevic, Alison E. Mather, John Wain, Mark Webber, Andrew J. Page, Justin O'Grady |
| EPI_ISL_1315160 | The Jackson Laboratory | The Jackson Laboratory | Bergeron D, Renzette N, Adams M, Omerza G, Kelly K, Li L |
| EPI_ISL_1323808 | National Virus Reference Laboratory | National Virus Reference Laboratory | Fiona Crispie, Calum Walsh, Zoe Yandle, Charlene Bennet, Gabriel Gonzalez, Michael Carr, Jonathan Dean, Paul Cotter, Cillian F De Gascun |
| EPI_ISL_1325124 | Lighthouse Lab in Milton Keynes | Wellcome Sanger Institute for the COVID-19 Genomics UK (COG-UK) Consortium | The Lighthouse Lab in Milton Keynes and Alex Alderton, Roberto Amato, Jeffrey Barrett, Sonia Goncalves, Ewan Harrison, David K. Jackson, Ian Johnston, Dominic Kwiatkowski, Cordelia Langford, John Sillitoe on behalf of the Wellcome Sanger Institute COVID-19 Surveillance Team |
| EPI_ISL_1327536 | Lighthouse Lab in Cambridge | Wellcome Sanger Institute for the COVID-19 Genomics UK (COG-UK) Consortium | Rob Howes, The Lighthouse Lab in Cambridge and Alex Alderton, Roberto Amato, Jeffrey Barrett, Sonia Goncalves, Ewan Harrison, David K. Jackson, Ian Johnston, Dominic Kwiatkowski, Cordelia Langford, John Sillitoe on behalf of the Wellcome Sanger Institute COVID-19 Surveillance Team |
| EPI_ISL_1333216, EPI_ISL_1333236 | Lighthouse Lab in Glasgow | Wellcome Sanger Institute for the COVID-19 Genomics UK (COG-UK) Consortium | Harper VanSteenhouse, Yumi Kasai, David Gray, Carol Clugston, Anna Dominiczak and Alex Alderton, Roberto Amato, Jeffrey Barrett, Sonia Goncalves, Ewan Harrison, David K. Jackson, Ian Johnston, Dominic Kwiatkowski, Cordelia Langford, John Sillitoe on behalf of the Wellcome Sanger Institute COVID-19 Surveillance Team |
| EPI_ISL_1333819, EPI_ISL_1333966 | Lighthouse Lab in Cambridge | Wellcome Sanger Institute for the COVID-19 Genomics UK (COG-UK) Consortium | Rob Howes, The Lighthouse Lab in Cambridge and Alex Alderton, Roberto Amato, Jeffrey Barrett, Sonia Goncalves, Ewan Harrison, David K. Jackson, Ian Johnston, Dominic Kwiatkowski, Cordelia Langford, John Sillitoe on behalf of the Wellcome Sanger Institute COVID-19 Surveillance Team |
| EPI_ISL_1334058, EPI_ISL_1334074 | Lighthouse Lab in Glasgow | Wellcome Sanger Institute for the COVID-19 Genomics UK (COG-UK) Consortium | Harper VanSteenhouse, Yumi Kasai, David Gray, Carol Clugston, Anna Dominiczak and Alex Alderton, Roberto Amato, Jeffrey Barrett, Sonia Goncalves, Ewan Harrison, David K. Jackson, Ian Johnston, Dominic Kwiatkowski, Cordelia Langford, John Sillitoe on behalf of the Wellcome Sanger Institute COVID-19 Surveillance Team |
| EPI_ISL_1340956 | 1. Glowny Inspektorat Sanitarny; 2. Diagnostyka. Laboratoria Medyczne. | 1. ViroGenetics - BSL3 Laboratory of Virology, Maopolska Centre of Biotechnology, Jagiellonian University; 2. Diagtron Laboratoria Lukasz Rabalski | Rabalski L., Gromowski,T., Mazur-Panasiuk,N., Kowalski,M., Maciej Kosinski, Natalia Derewonko, Szulc,P., Sylwia Januszczak, Labaj,P.P., Pyrc,K. |
| EPI_ISL_1342703 | Lighthouse Lab in Alderley Park | Wellcome Sanger Institute for the COVID-19 Genomics UK (COG-UK) Consortium | Jacquelyn Wynn, Mairead Hyland, The Lighthouse Lab in Alderley Park and Alex Alderton, Roberto Amato, Jeffrey Barrett, Sonia Goncalves, Ewan Harrison, David K. Jackson, Ian Johnston, Dominic Kwiatkowski, Cordelia Langford, John Sillitoe on behalf of the Wellcome Sanger Institute COVID-19 Surveillance Team |
| EPI_ISL_1381817 | Hospital | National Reference Center for Viruses of Respiratory Infections, Institut Pasteur, Paris | Marion Barbet, Sylvie Behillil, Méline Bizard, Angela Brisebarre, Camille Capel, Louise Lefrançois, Etienne Simon-Lorière, Vincent Enouf, Maud Vanpeene, Sylvie van der Werf,Leruez-Ville Marianne |
| EPI_ISL_1385215 | Pandemic Response Lab - NYC | Pandemic Response Lab, R&D | Henry Lee, Michael Hammerling, Melissa Hopkins, Cybill del Castillo, Shinyoung Clair Kang, William Ward, Pradeep Bugga, Sol Rey, Dylan Law, Haiping Hao, Jon Laurent |
| EPI_ISL_1389320 | Labo Analyses Med | National Reference Center for Viruses of Respiratory Infections, Institut Pasteur, Paris | Marion Barbet, Sylvie Behillil, Méline Bizard, Angela Brisebarre, Camille Capel, Louise Lefrançois, Etienne Simon-Lorière, Vincent Enouf, Maud Vanpeene, Sylvie van der Werf,Felloni Claire |
| EPI_ISL_1397670, EPI_ISL_1397673 | NORTH SHORE UNIVERSITY HOSPITAL | Wadsworth Center, New York State Department of Health | Kirsten St. George, Daryl M. Lamson, Alexis Russel, Matthew Shudt, Melissa A Leisner, Jonathan Plitnick, Navjot Singh, John Kelly, Erasmus Schneider, Erica Lasek-Nesselquist |
| EPI_ISL_1404435 | Università di Parma, Laboratorio di Igiene e Sanità Pubblica | Istituto Zooprofilattico Sperimentale della Lombardia e dell'Emilia Romagna (IZSLER), Risk Analysis and Genomic Epidemiology Unit | Maria Eugenia Colucci, Licia Veronesi, Paola Affanni, Marina Morganti, Ilaria Menozzi, Erika Scaltriti, Stefano Pongolini |
| EPI_ISL_1404459 | Hôpital Necker-Enfants malades | Department of Virology, Henri Mondor University Hospital, Assistance Publique Hôpitaux de Paris, Université Paris-Est Créteil, INSERM U955 | Christophe Rodriguez, Slim Fourati, Vanessa Demontant, Guillaume Gricourt, Melissa N'Debi, Alexandre Soulier, Elisabeth Trawinski, Jean-Michel Pawlotsky |
| EPI_ISL_1410170 | Lighthouse Lab in Cambridge | Wellcome Sanger Institute for the COVID-19 Genomics UK (COG-UK) Consortium | Rob Howes, The Lighthouse Lab in Cambridge and Alex Alderton, Roberto Amato, Jeffrey Barrett, Sonia Goncalves, Ewan Harrison, David K. Jackson, Ian Johnston, Dominic Kwiatkowski, Cordelia Langford, John Sillitoe on behalf of the Wellcome Sanger Institute COVID-19 Surveillance Team |
| EPI_ISL_1410622 | Lighthouse Lab in Alderley Park | Wellcome Sanger Institute for the COVID-19 Genomics UK (COG-UK) Consortium | Jacquelyn Wynn, Mairead Hyland, The Lighthouse Lab in Alderley Park and Alex Alderton, Roberto Amato, Jeffrey Barrett, Sonia Goncalves, Ewan Harrison, David K. Jackson, Ian Johnston, Dominic Kwiatkowski, Cordelia Langford, John Sillitoe on behalf of the Wellcome Sanger Institute COVID-19 Surveillance Team |
| EPI_ISL_1411717, EPI_ISL_1411771, | Lighthouse Lab in Milton Keynes | Wellcome Sanger Institute for the COVID-19 Genomics UK | The Lighthouse Lab in Milton Keynes and Alex Alderton, Roberto Amato, Jeffrey Barrett, Sonia Goncalves, Ewan Harrison, David K. Jackson, Ian |

| | | | |
|---|---|---|---|
| EPI_ISL_1415939, EPI_ISL_1415951 | | (COG-UK) Consortium | Johnston, Dominic Kwiatkowski, Cordelia Langford, John Sillitoe on behalf of the Wellcome Sanger Institute COVID-19 Surveillance Team |
| EPI_ISL_1417738, EPI_ISL_1417975 | Swedish national genomic surveillance program of SARS-CoV-2 | The Public Health Agency of Sweden | Swedish national genomic surveillance program of SARS-CoV-2 |
| EPI_ISL_1421767, EPI_ISL_1422066 | Laboratory Corporation of America | Centers for Disease Control and Prevention Division of Viral Diseases, Pathogen Discovery | Peter W. Cook, Dakota Howard, Dhwani Batra, Ben L. Rambo-Martin, Minoo Agarwal, Eyad Almasri, Debbie Boles, Ayla Burns, Nuthawin Charoensri, Oren Cohen, Susan Countryman, Mary Ann Cristobal, Bobbi Croy, Suzanne Dale, Hrushikesh Deshmukh, Amanda Douglas, Vincent Drouillon, Marcia Eisenberg, Howard Engler, Rama Ghatti, Prashant Gupta, Susan Hicks, Jake Humphrey, Lax Iyer, Manoj Jain, Mohan Kolli, Brian Krueger, Tim Kuphal, Stanley Letovsky, Michael Levandoski, Craig Lukasik, Jonathan Meltzer, Brian Norvell, Mindy Nye, Scott Parker, Christos Petropoulos, John Pruitt, Steven Ragan, Scott Ryan, Mike Sapeta, Jana Schroth, Suresh Babu Selvaraju, Goran Stevovic, Amanda Suchanek, Andrea Throop, Lyndon Tilson, Thomas Urban, Joe Voshell, Kimberly Wagner, Jonathan Williams, Mary Williamson, Qian Zeng, Tricia Zwiefelhofer, Clinton R. Paden, Suxiang Tong, Duncan MacCannell |
| EPI_ISL_1437372 | SYNLAB MVZ Leverkusen | Robert Koch Institute | unknown |
| EPI_ISL_1439178 | LabKom - MVZ Labor Bochum MLB GmbH | Robert Koch Institute | unknown |
| EPI_ISL_1444902 | Maryland Genomics, Institute for Genome Sciences, University of Maryland School of Medicine | Maryland Genomics, Institute for Genome Sciences, University of Maryland School of Medicine | Tallon, Luke J; Sadzewicz, Lisa D; Humphrys, Mike; Ott, Sandra; Roussey, Holly; Mehta, Aditya; Vavikolanu, Kranthi; Fraser, Claire M; Ravel, Jacques |
| EPI_ISL_1447794 | IZSM | TIGEM | Antonio Grimaldi Patrizia Annunziata Francesco Panariello Biancamaria Pierri Claudia Tiberio Valentina Bouche Chiara Colantuono Maria Concetta Cuomo Denise Di Concilio Lucio Di Filippo Anna Manfredi Marcello Salvi Antonio Limone Luigi Atripaldi Pellegrino Cerino Andrea Ballabio Davide Cacchiarelli |
| EPI_ISL_1455912 | Dutch COVID-19 response team | National Institute for Public Health and the Environment (RIVM) | Adam Meijer, Harry Vennema, Dirk Eggink, Jeroen Cremer, Sharon van den Brink, Bas van der Veer, AnneMarie van den Brandt, Lisa Wijsman, Kim Freriks, Ryanne Jaarsma, Eunìce Then, Jolienke Hardeman, Lynn Aarts, Sanne Bos, Melissa van Tuil, Robert Kohl, Linda van de Nes, Sjoerd Kuiling, James Groot, Florian Zwagemaker, Dennis Schmitz, Annelies Kroneman, Karim Hajji, Chantal Reusken, on behalf of the national COVID-19 response team |
| EPI_ISL_1468017 | Lighthouse Lab in Milton Keynes | Wellcome Sanger Institute for the COVID-19 Genomics UK (COG-UK) Consortium | The Lighthouse Lab in Milton Keynes and Alex Alderton, Roberto Amato, Jeffrey Barrett, Sonia Goncalves, Ewan Harrison, David K. Jackson, Ian Johnston, Dominic Kwiatkowski, Cordelia Langford, John Sillitoe on behalf of the Wellcome Sanger Institute COVID-19 Surveillance Team |
| EPI_ISL_1468482 | Clinical Microbiology, Infection Prevention and Control | Section for Molecular Diagnostics | Björn Hallström, Jonas Björkman |
| EPI_ISL_1480225 | Helix/Illumina | Centers for Disease Control and Prevention Division of Viral Diseases, Pathogen Discovery | Dakota Howard, Dhwani Batra, Peter W. Cook, Kara Moser, Adrian Paskey, Jason Caravas, Benjamin Rambo-Martin, Shatavia Morrison, Christopher Gulvick, Scott Sammons, Yvette Unoarumhi, Darlene Wagner, Matthew Schmerer, Eileen de Feo, Jan Antico, Christine Tran, Matthew Tolentino, Shannon Wickline, Kim Gietzen, Brad Sickler, Jingtao Liu, Eric Allen, Phil Febbo, Nicole L. Washington, Simon White, Geraint Levan, Kelly Schiabor Barrett, Elizabeth Cirulli, Alexandre Bolze, Ary Ascencio, Charlotte Rivera-Garcia, Ryan Cho, Jason Nguyen, Sherry Wang, Jimmy Ramirez, Tyler Cassens, Efren Sandoval, Magnus Isaksson, William Lee, David Becker, Marc Laurent, James Lu, Clinton R. Paden, Duncan MacCannell |
| EPI_ISL_1486118 | Lighthouse Lab in Glasgow | Wellcome Sanger Institute for the COVID-19 Genomics UK (COG-UK) Consortium | Harper VanSteenhouse, Yumi Kasai, David Gray, Carol Clugston, Anna Dominiczak and Alex Alderton, Roberto Amato, Jeffrey Barrett, Sonia Goncalves, Ewan Harrison, David K. Jackson, Ian Johnston, Dominic Kwiatkowski, Cordelia Langford, John Sillitoe on behalf of the Wellcome Sanger Institute COVID-19 Surveillance Team |
| EPI_ISL_1487655 | Lighthouse Lab in Milton Keynes | Wellcome Sanger Institute for the COVID-19 Genomics UK (COG-UK) Consortium | The Lighthouse Lab in Milton Keynes and Alex Alderton, Roberto Amato, Jeffrey Barrett, Sonia Goncalves, Ewan Harrison, David K. Jackson, Ian Johnston, Dominic Kwiatkowski, Cordelia Langford, John Sillitoe on behalf of the Wellcome Sanger Institute COVID-19 Surveillance Team |
| EPI_ISL_1488119 | Lighthouse Lab in Alderley Park | Wellcome Sanger Institute for the COVID-19 Genomics UK (COG-UK) Consortium | Jacquelyn Wynn, Mairead Hyland, The Lighthouse Lab in Alderley Park and Alex Alderton, Roberto Amato, Jeffrey Barrett, Sonia Goncalves, Ewan Harrison, David K. Jackson, Ian Johnston, Dominic Kwiatkowski, Cordelia Langford, John Sillitoe on behalf of the Wellcome Sanger Institute COVID-19 Surveillance Team |
| EPI_ISL_1497724, EPI_ISL_1499206 | 1. Glowny Inspektorat Sanitarny. 2. Diagnostyka. Laboratoria Medyczne. | 1. ViroGenetics - BSL3 Laboratory of Virology, Maopolska Centre of Biotechnology, Jagiellonian University; 2. genXone SA, Research & Development Laboratory | Mazur-Panasiuk,N., Grzegorz Nowicki, Gromowski,T., Natalia Drweska-Matelska, Jakub Grabowski, Anna Brylak, Aleksandra Gidlewicz, Karol Szeszko, Maciej Sykulski, ukasz Krych, Kowalski,M., Szulc,P., Sylwia Januszczak, Labaj,P.P., Micha Kaszuba, Pyrc,K. |
| EPI_ISL_1504129, EPI_ISL_1504295 | Lighthouse Lab in Alderley Park | Wellcome Sanger Institute for the COVID-19 Genomics UK (COG-UK) Consortium | Jacquelyn Wynn, Mairead Hyland, The Lighthouse Lab in Alderley Park and Alex Alderton, Roberto Amato, Jeffrey Barrett, Sonia Goncalves, Ewan Harrison, David K. Jackson, Ian Johnston, Dominic Kwiatkowski, Cordelia Langford, John Sillitoe on behalf of the Wellcome Sanger Institute COVID-19 Surveillance Team |
| EPI_ISL_1517094 | Armies | National Reference Center for Viruses of Respiratory Infections, Institut Pasteur, Paris | Marion Barbet, Sylvie Behillil, Méline Bizard, Angela Brisebarre, Camille Capel,Frédéric Lemoine,Corinne Maufrais,Christophe Malabat, Louise Lefrançois, Etienne Simon-Lorière, Vincent Enouf, Maud Vanpeene, Sylvie van der Werf,Marine Desroches |
| EPI_ISL_1517226, EPI_ISL_1517227 | Labo Analyses Med | National Reference Center for Viruses of Respiratory Infections, Institut Pasteur, Paris | Marion Barbet, Sylvie Behillil, Méline Bizard, Angela Brisebarre, Camille Capel,Frédéric Lemoine,Corinne Maufrais,Christophe Malabat, Louise Lefrançois, Etienne Simon-Lorière, Vincent Enouf, Maud Vanpeene, Sylvie van der Werf,Nabil Gastli |
| EPI_ISL_1531761 | University of Oregon COVID-19 MAP Laboratory | University of Oregon Genomics and Cell Characterization Core Facility (GC3F) | Douglas Turnbull, Ariana White, Jeff Bishop, Jason Carriere |
| EPI_ISL_1534045, EPI_ISL_1534092 | Clinical Microbiology, Infection Prevention and Control | Section for Molecular Diagnostics | Björn Hallström, Jonas Björkman |
| EPI_ISL_1534527 | Ministry of Health Turkey | Ministry of Health Turkey | Fatma Bayrakdar, Yasemin Cosgun, Suleyman Yalcin, Gulay Korukluoglu |
| EPI_ISL_1535235, EPI_ISL_1535394, EPI_ISL_1535527 | Lighthouse Lab in Milton Keynes | Wellcome Sanger Institute for the COVID-19 Genomics UK (COG-UK) Consortium | The Lighthouse Lab in Milton Keynes and Alex Alderton, Roberto Amato, Jeffrey Barrett, Sonia Goncalves, Ewan Harrison, David K. Jackson, Ian Johnston, Dominic Kwiatkowski, Cordelia Langford, John Sillitoe on behalf of the Wellcome Sanger Institute COVID-19 Surveillance Team |
| EPI_ISL_1535641, EPI_ISL_1535771, EPI_ISL_1536308, EPI_ISL_1536532 | Lighthouse Lab in Alderley Park | Wellcome Sanger Institute for the COVID-19 Genomics UK (COG-UK) Consortium | Jacquelyn Wynn, Mairead Hyland, The Lighthouse Lab in Alderley Park and Alex Alderton, Roberto Amato, Jeffrey Barrett, Sonia Goncalves, Ewan Harrison, David K. Jackson, Ian Johnston, Dominic Kwiatkowski, Cordelia Langford, John Sillitoe on behalf of the Wellcome Sanger Institute COVID-19 Surveillance Team |
| EPI_ISL_1536669 | Lighthouse Lab in Milton Keynes | Wellcome Sanger Institute for the COVID-19 Genomics UK (COG-UK) Consortium | The Lighthouse Lab in Milton Keynes and Alex Alderton, Roberto Amato, Jeffrey Barrett, Sonia Goncalves, Ewan Harrison, David K. Jackson, Ian Johnston, Dominic Kwiatkowski, Cordelia Langford, John Sillitoe on behalf of the Wellcome Sanger Institute COVID-19 Surveillance Team |
| EPI_ISL_1536850, EPI_ISL_1537040 | Lighthouse Lab in Alderley Park | Wellcome Sanger Institute for the COVID-19 Genomics UK (COG-UK) Consortium | Jacquelyn Wynn, Mairead Hyland, The Lighthouse Lab in Alderley Park and Alex Alderton, Roberto Amato, Jeffrey Barrett, Sonia Goncalves, Ewan Harrison, David K. Jackson, Ian Johnston, Dominic Kwiatkowski, Cordelia Langford, John Sillitoe on behalf of the Wellcome Sanger Institute COVID-19 Surveillance Team |
| EPI_ISL_1542234 | Pandemic Response Lab - NYC | Pandemic Response Lab, R&D | Henry Lee, Michael Hammerling, Melissa Hopkins, Cybill del Castillo, Shinyoung Clair Kang, William Ward, Pradeep Bugga, Sol Rey, Dylan Law, Katharine Nelson, Haiping Hao, Jon Laurent |
| EPI_ISL_1545475 | Istituto Zooprofilattico Sperimentale del Mezzogiorno - Azienda Ospedaliera Pugliese Ciaccio di Catanzaro | TIGEM | Antonio Grimaldi Patrizia Annunziata Francesco Panariello Biancamaria Pierri Claudia Tiberio Teresa Giuliano Valentina Bouche Chiara Colantuono Maria Concetta Cuomo Denise Di Concilio Lucio Di Filippo Anna Manfredi Marcello Salvi Pasquale Minchella Antonio Limone Luigi Atripaldi Pellegrino Cerino Andrea Ballabio Davide Cacchiarelli |
| EPI_ISL_1545801 | University of Exeter | COVID-19 Genomics UK (COG-UK) Consortium | Ben Temperton,Aaron Jeffries,Michelle Michelsen,Joanna Warwick-Dugdale,Audrey Farbos,Robyn Manley,Stephen Michell,Jane Masoli |
| EPI_ISL_1546148, EPI_ISL_1546187 | Liverpool Clinical Laboratories | COVID-19 Genomics UK (COG-UK) Consortium | Sam Haldenby, Alistair Darby, Steve Paterson, Anita Lucaci, Julian Hiscox, M Almsaud, A Alrezaihi, Muhannad Alruwaili, Stuart D Armstrong, Jones Benjamin, Eleanor G Bentley, Anu Chawla, Jordan J Clark, Angela Cowell, Richard Eccles, Isabel García-Dorival, Matthew Gemmell, Alessandro Gerada, PKF Gilmore, Richard Gregory, Ximeng Han, Catherine Hartley, Margaret Hughes, Miren Iturriza-Gomara, James Johnson, L Luu, Jenifer Manson, Charlotte Nelson, Elaine O'Toole, Cassie Olateju, Rebekah Penrice-Randal , Lucille Rainbow, N.P Randle, Trevor Ian Robinson, Parul Sharma, Ghada T Shawli, James P Stewart, Neil Swainston, Ecaterina Vamos, Joanne Watts, Mark Whitehead, Hermione Webster |
| EPI_ISL_1548232, EPI_ISL_1548269, EPI_ISL_1548379, EPI_ISL_1548676, | Laboratory Corporation of America | Centers for Disease Control and Prevention Division of Viral Diseases, Pathogen Discovery | Dakota Howard, Dhwani Batra, Peter W. Cook, Kara Moser, Adrian Paskey, Jason Caravas, Benjamin Rambo-Martin, Shatavia Morrison, Christopher Gulvick, Scott Sammons, Yvette Unoarumhi, Darlene Wagner, Matthew Schmerer, Minoo Agarwal, Eyad Almasri, Debbie Boles, Ayla Burns, Nuthawin |

| | | | |
|---|---|---|---|
| EPI_ISL_1548767, EPI_ISL_1549960 | | | Charoensri, Oren Cohen, Susan Countryman, Mary Ann Cristobal, Bobbi Croy, Suzanne Dale, Hrushikesh Deshmukh, Amanda Douglas, Vincent Drouillon, Marcia Eisenberg, Howard Engler, Rama Ghatti, Prashant Gupta, Susan Hicks, Jake Humphrey, Lax Iyer, Manoj Jain, Mohan Kolli, Brian Krueger, Tim Kuphal, Stanley Letovsky, Michael Levandoski, Craig Lukasik, Jonathan Meltzer, Brian Norvell, Mindy Nye, Scott Parker, Christos Petropoulos, John Pruitt, Steven Ragan, Scott Ryan, Mike Sapeta, Jana Schroth, Suresh Babu Selvaraju, Goran Stevovic, Amanda Suchanek, Andrea Throop, Lyndon Tilson, Thomas Urban, Joe Voshell, Kimberly Wagner, Jonathan Williams, Mary Williamson, Qian Zeng, Tricia Zwiefelhofer, Clinton R. Paden, Duncan MacCannell |
| EPI_ISL_1552460 | Quest Diagnostics Incorporated | Centers for Disease Control and Prevention Division of Viral Diseases, Pathogen Discovery | Dakota Howard, Dhwani Batra, Peter W. Cook, Kara Moser, Adrian Paskey, Jason Caravas, Benjamin Rambo-Martin, Shatavia Morrison, Christopher Gulvick, Scott Sammons, Yvette Unoarumhi, Darlene Wagner, Matthew Schmerer, S. H. Rosenthal, A. Gerasimova, R. M. Kagan, B. Anderson, M. Hua, Y. Liu, L.E. Bernstein, K.E. Livingston, A. Perez, I. A. Shlyakhter, R. V. Rolando, R. Owen, P. Tanpaiboon, F. Lacbawan, Clinton R. Paden, Duncan MacCannell |
| EPI_ISL_1553367, EPI_ISL_1554137, EPI_ISL_1554345, EPI_ISL_1554495, EPI_ISL_1554512 | Helix/Illumina | Centers for Disease Control and Prevention Division of Viral Diseases, Pathogen Discovery | Dakota Howard, Dhwani Batra, Peter W. Cook, Kara Moser, Adrian Paskey, Jason Caravas, Benjamin Rambo-Martin, Shatavia Morrison, Christopher Gulvick, Scott Sammons, Yvette Unoarumhi, Darlene Wagner, Matthew Schmerer, Eileen de Feo, Jan Antico, Christine Tran, Matthew Tolentino, Shannon Wickline, Kim Gietzen, Brad Sickler, Jingtao Liu, Eric Allen, Phil Febbo, Nicole L. Washington, Simon White, Geraint Levan, Kelly Schiabor Barrett, Elizabeth Cirulli, Alexandre Bolze, Ary Ascencio, Charlotte Rivera-Garcia, Ryan Cho, Jason Nguyen, Sherry Wang, Jimmy Ramirez, Tyler Cassens, Efren Sandoval, Magnus Isaksson, William Lee, David Becker, Marc Laurent, James Lu, Clinton R. Paden, Duncan MacCannell |
| EPI_ISL_1555203, EPI_ISL_1555522, EPI_ISL_1555559, EPI_ISL_1556044, EPI_ISL_1556270 | Fulgent Genetics | Centers for Disease Control and Prevention Division of Viral Diseases, Pathogen Discovery | Dakota Howard, Dhwani Batra, Peter W. Cook, Kara Moser, Adrian Paskey, Jason Caravas, Benjamin Rambo-Martin, Shatavia Morrison, Christopher Gulvick, Scott Sammons, Yvette Unoarumhi, Darlene Wagner, Matthew Schmerer, Harry Gao, Mickey Li, John Gao, Joseph Fierro, Benafsh Sapra, Becky Tsai, Yan Meng, Doreen Ng, James Xie, Clinton R. Paden, Duncan MacCannell |
| EPI_ISL_1558386 | ASL Napoli 1 Centro | AMES Centro Polidiagnostico Strumentale S.r.l. | "Giovanni Savarese, Raffaella Ruggiero, Eloisa Evangelista, Antonella Di Carlo, Luisa Circelli, Luigi D'Amore, Nadia Petrillo, Monica Ianniello, Roberto Sirica,Maurizio D'Amora, Antonio Fico" |
| EPI_ISL_1561041, EPI_ISL_1562092, EPI_ISL_1563386 | Aegis Sciences Corporation | Centers for Disease Control and Prevention Division of Viral Diseases, Pathogen Discovery | Dakota Howard, Dhwani Batra, Peter W. Cook, Kara Moser, Adrian Paskey, Jason Caravas, Benjamin Rambo-Martin, Shatavia Morrison, Christopher Gulvick, Scott Sammons, Yvette Unoarumhi, Darlene Wagner, Matthew Schmerer, Cyndi Clark, Patrick Campbell, Rob Case, Vikramsinha Ghorpade, Holly Houdeshell, Ola Kvalvaag, Dillon Nall, Ethan Sanders, Alec Vest, Shaun Westlund, Matthew Hardison, Clinton R. Paden, Duncan MacCannell |
| EPI_ISL_1563971 | Virologisches Institut des UniversitÃ¤tsklinikums Erlangen | Robert Koch Institute | unknown |
| EPI_ISL_1564551, EPI_ISL_1564656 | Lighthouse Lab in Alderley Park | Wellcome Sanger Institute for the COVID-19 Genomics UK (COG-UK) Consortium | Jacquelyn Wynn, Mairead Hyland, The Lighthouse Lab in Alderley Park and Alex Alderton, Roberto Amato, Jeffrey Barrett, Sonia Goncalves, Ewan Harrison, David K. Jackson, Ian Johnston, Dominic Kwiatkowski, Cordelia Langford, John Sillitoe on behalf of the Wellcome Sanger Institute COVID-19 Surveillance Team |
| EPI_ISL_1565471 | IMD - MVZ Labor Martinsried | Robert Koch Institute | unknown |
| EPI_ISL_1566469, EPI_ISL_1566487 | Synlab MVZ Augsburg | Robert Koch Institute | unknown |
| EPI_ISL_1568108 | Dianovis GmbH Greiz | Robert Koch Institute | unknown |
| EPI_ISL_1570373 | Bayerisches Landesamt fÃ¼r Gesundheit und Lebensmittelsicherheit (LGL) | Robert Koch Institute | unknown |
| EPI_ISL_1571623 | SYNLAB Labor MÃ¼nchen Zentrum LMZ | Robert Koch Institute | unknown |
| EPI_ISL_1572035, EPI_ISL_1572037 | Klinikum der Stadt Ludwigshafen - Institut fÃ¼r Labordiagnostik Hygiene und Transfusionsmedizin | Robert Koch Institute | unknown |
| EPI_ISL_1574410, EPI_ISL_1574502, EPI_ISL_1574635 | SYNLAB MVZ Weiden | Robert Koch Institute | unknown |
| EPI_ISL_1574650 | SYNLAB MVZ Dachau | Robert Koch Institute | unknown |
| EPI_ISL_1575645, EPI_ISL_1575849, EPI_ISL_1576144, EPI_ISL_1581406 | Helix/Illumina | Centers for Disease Control and Prevention Division of Viral Diseases, Pathogen Discovery | Dakota Howard, Dhwani Batra, Peter W. Cook, Kara Moser, Adrian Paskey, Jason Caravas, Benjamin Rambo-Martin, Shatavia Morrison, Christopher Gulvick, Scott Sammons, Yvette Unoarumhi, Darlene Wagner, Matthew Schmerer, Eileen de Feo, Jan Antico, Christine Tran, Matthew Tolentino, Shannon Wickline, Kim Gietzen, Brad Sickler, Jingtao Liu, Eric Allen, Phil Febbo, Nicole L. Washington, Simon White, Geraint Levan, Kelly Schiabor Barrett, Elizabeth Cirulli, Alexandre Bolze, Ary Ascencio, Charlotte Rivera-Garcia, Ryan Cho, Jason Nguyen, Sherry Wang, Jimmy Ramirez, Tyler Cassens, Efren Sandoval, Magnus Isaksson, William Lee, David Becker, Marc Laurent, James Lu, Clinton R. Paden, Duncan MacCannell |
| EPI_ISL_1582032, EPI_ISL_1582107 | Quest Diagnostics Incorporated | Centers for Disease Control and Prevention Division of Viral Diseases, Pathogen Discovery | Dakota Howard, Dhwani Batra, Peter W. Cook, Kara Moser, Adrian Paskey, Jason Caravas, Benjamin Rambo-Martin, Shatavia Morrison, Christopher Gulvick, Scott Sammons, Yvette Unoarumhi, Darlene Wagner, Matthew Schmerer, S. H. Rosenthal, A. Gerasimova, R. M. Kagan, B. Anderson, M. Hua, Y. Liu, L.E. Bernstein, K.E. Livingston, A. Perez, I. A. Shlyakhter, R. V. Rolando, R. Owen, P. Tanpaiboon, F. Lacbawan, Clinton R. Paden, Duncan MacCannell |
| EPI_ISL_1583042 | Hospital | National Reference Center for Viruses of Respiratory Infections, Institut Pasteur, Paris | Marion Barbet, Sylvie Behillil, Frédéric Lemoine, Corinne Maufrais, Christophe Malabat, Méline Bizard, Angela Brisebarre, Camille Capel, Louise Lefrançois, Etienne Simon-Lorière, Vincent Enouf, Maud Vanpeene, Sylvie van der Werf,LéA Pilorge |
| EPI_ISL_1583447 | Department of Microbiology, University Innsbruck | Bergthaler laboratory, CeMM Research Center for Molecular Medicine of the Austrian Academy of Sciences | Lukas Endler, Anna Schedl, Fabian Amman, Petr Triska, Thomas Penz, Benedikt Agerer, Maelle Le Moing, Michael Schuster, Bekir Erguner, Jan Laine, Martin Senekowitsch, Christoph Bock, Andreas Bergthaler |
| EPI_ISL_1583457 | Elling group, Institute of Molecular Biotechnology (IMBA) | Bergthaler laboratory, CeMM Research Center for Molecular Medicine of the Austrian Academy of Sciences | Lukas Endler, Anna Schedl, Fabian Amman, Petr Triska, Thomas Penz, Benedikt Agerer, Maelle Le Moing, Michael Schuster, Bekir Erguner, Jan Laine, Martin Senekowitsch, Christoph Bock, Andreas Bergthaler |
| EPI_ISL_1585913 | University of Michigan Clinical Microbiology Laboratory | Lauring Lab, University of Michigan, Department of Microbiology and Immunology | Valesano |
| EPI_ISL_1592444, EPI_ISL_1592565 | Helix/Illumina | Centers for Disease Control and Prevention Division of Viral Diseases, Pathogen Discovery | Dakota Howard, Dhwani Batra, Peter W. Cook, Kara Moser, Adrian Paskey, Jason Caravas, Benjamin Rambo-Martin, Shatavia Morrison, Christopher Gulvick, Scott Sammons, Yvette Unoarumhi, Darlene Wagner, Matthew Schmerer, Eileen de Feo, Jan Antico, Christine Tran, Matthew Tolentino, Shannon Wickline, Kim Gietzen, Brad Sickler, Jingtao Liu, Eric Allen, Phil Febbo, Nicole L. Washington, Simon White, Geraint Levan, Kelly Schiabor Barrett, Elizabeth Cirulli, Alexandre Bolze, Ary Ascencio, Charlotte Rivera-Garcia, Ryan Cho, Jason Nguyen, Sherry Wang, Jimmy Ramirez, Tyler Cassens, Efren Sandoval, Magnus Isaksson, William Lee, David Becker, Marc Laurent, James Lu, Clinton R. Paden, Duncan MacCannell |
| EPI_ISL_1593839 | National Institute of Public Health | State Veterinary Institute Prague | Nagy,A;Jirincova,H;Suri,T;Trnka,D;Vecerova,J |
| EPI_ISL_1594447 | Labo Analyses Med | National Reference Center for Viruses of Respiratory Infections, Institut Pasteur, Paris | Marion Barbet, Sylvie Behillil, Frédéric Lemoine, Corinne Maufrais, Christophe Malabat, Méline Bizard, Angela Brisebarre, Camille Capel, Louise Lefrançois, Etienne Simon-Lorière, Vincent Enouf, Maud Vanpeene, Sylvie van der Werf,Pierre-Yves Leonard |
| EPI_ISL_1596007, EPI_ISL_1596882, EPI_ISL_1596917 | Dutch COVID-19 response team | National Institute for Public Health and the Environment (RIVM) | Adam Meijer, Harry Vennema, Dirk Eggink, Jeroen Cremer, Sharon van den Brink, Bas van der Veer, AnneMarie van den Brandt, Lisa Wijsman, Kim Freriks, Ryanne Jaarsma, Euníce Then, Jolienke Hardeman, Lynn Aarts, Sanne Bos, Melissa van Tuil, Robert Kohl, Linda van de Nes, Sjoerd Kuiling, James Groot, Florian Zwagemaker, Dennis Schmitz, Annelies Kroneman, Karim Hajji, Chantal Reusken, on behalf of the national COVID-19 response team |
| EPI_ISL_1598498 | Viollier AG | Department of Biosystems Science and Engineering, ETH Zürich | Christian Beisel, Sarah Nadeau, Chaoran Chen, Ivan Topolsky, Philipp Jablonski, Lara Fuhrmann, David Dreifuss, Katharina Jahn, Rebecca Denes, Mirjam Feldkamp, Ina Nissen, Natascha Santacroce, Elodie Burcklen, Christiane Beckmann, Maurice Redondo, Olivier Kobel, Christoph Noppen, Sophie Seidel, Noemie Santamaria de Souza, Niko Beerenwinkel, Tanja Stadler |
| EPI_ISL_1599263, EPI_ISL_1599264 | Public Health Authority of the Slovak Republic | Laboratory of Genomics and Bioinformatics, Comenius University Science Park | Tatiana Sedláková, Diana Rusáková, Miroslav Böhmer, Anna Giová, Jaroslav Budiš, Tomáš Szemes |
| EPI_ISL_1603155, EPI_ISL_1603909, EPI_ISL_1604246, EPI_ISL_1604766, EPI_ISL_1605457, EPI_ISL_1605460, EPI_ISL_1605765, EPI_ISL_1606419, EPI_ISL_1607160, EPI_ISL_1607184, EPI_ISL_1607361, EPI_ISL_1607422, EPI_ISL_1607439, EPI_ISL_1607483, EPI_ISL_1607487 | | | |
| see above | Swedish national genomic surveillance program of | The Public Health Agency of Sweden | Swedish national genomic surveillance program of SARS-CoV-2 |

| | | | |
|---|---|---|---|
| | SARS-CoV-2 | | |
| EPI_ISL_1608677 | Maryland Genomics, Institute for Genome Sciences, University of Maryland School of Medicine | Maryland Genomics, Institute for Genome Sciences, University of Maryland School of Medicine | Tallon, Luke J; Sadzewicz, Lisa D; Humphrys, Mike; Ott, Sandra; Roussey, Holly; Mehta, Aditya; Vavikolanu, Kranthi; Fraser, Claire M; Ravel, Jacques |
| EPI_ISL_1609898, EPI_ISL_1610474 | Laboratory Corporation of America | Centers for Disease Control and Prevention Division of Viral Diseases, Pathogen Discovery | Dakota Howard, Dhwani Batra, Peter W. Cook, Kara Moser, Adrian Paskey, Jason Caravas, Benjamin Rambo-Martin, Shatavia Morrison, Christopher Gulvick, Scott Sammons, Yvette Unoarumhi, Darlene Wagner, Matthew Schmerer, Minoo Agarwal, Eyad Almasri, Debbie Boles, Ayla Burns, Nuthawin Charoensri, Oren Cohen, Susan Countryman, Mary Ann Cristobal, Bobbi Croy, Suzanne Dale, Hrushikesh Deshmukh, Amanda Douglas, Vincent Drouillon, Marcia Eisenberg, Howard Engler, Rama Ghatti, Prashant Gupta, Susan Hicks, Jake Humphrey, Lax Iyer, Manoj Jain, Mohan Kolli, Brian Krueger, Tim Kuphal, Stanley Letovsky, Michael Levandoski, Craig Lukasik, Jonathan Meltzer, Brian Norvell, Mindy Nye, Scott Parker, Christos Petropoulos, John Pruitt, Steven Ragan, Scott Ryan, Mike Sapeta, Jana Schroth, Suresh Babu Selvaraju, Goran Stevovic, Amanda Suchanek, Andrea Throop, Lyndon Tilson, Thomas Urban, Joe Voshell, Kimberly Wagner, Jonathan Williams, Mary Williamson, Qian Zeng, Tricia Zwiefelhofer, Clinton R. Paden, Duncan MacCannell |
| EPI_ISL_1614046, EPI_ISL_1614384, EPI_ISL_1614834 | Fulgent Genetics | Centers for Disease Control and Prevention Division of Viral Diseases, Pathogen Discovery | Dakota Howard, Dhwani Batra, Peter W. Cook, Kara Moser, Adrian Paskey, Jason Caravas, Benjamin Rambo-Martin, Shatavia Morrison, Christopher Gulvick, Scott Sammons, Yvette Unoarumhi, Darlene Wagner, Matthew Schmerer, Harry Gao, Mickey Li, John Gao, Joseph Fierro, Benafsh Sapra, Becky Tsai, Yan Meng, Doreen Ng, James Xie, Clinton R. Paden, Duncan MacCannell |
| EPI_ISL_1616622 | UW Virology Lab | UW Virology Lab | Pavitra Roychoudhury, Hong Xie, Lasata Shrestha, Shah Mohamed Bakhash, Michelle Lin, Noah R. Baker, Sean Ellis, Saraswathi Sathees, Meei-Li Huang, Keith R Jerome, Alexander Greninger |
| EPI_ISL_1617248 | Colorado Department of Public Health and Environment | Colorado Department of Public Health and Environment | Laura Bankers, Molly C. Hetherington-Rauth, Diana Ir, Shannon Ely, Shannon R. Matzinger, Sarah Elizabeth Totten, Emily A. Travanty |
| EPI_ISL_1621352 | Hospital of the University of Pennsylvania Molecular Pathology Lab | Bushman Lab - University of Pennsylvania | John Everett, Kyle Rodino, Shantan Reddy, Pascha Hokama, Aoife M. Roche, Young Hwang, Abigail Glascock, Scott Sherrill-Mix, Samantha A. Whiteside, Jevon Graham-Wooten, Layla A. Khatib, Ayannah S. Fitzgerald, Arupa Ganguly, Mike Feldman, Brendan Kelly, Ronald G. Collman and Frederic Bushman |
| EPI_ISL_1621485 | Hospital | National Reference Center for Viruses of Respiratory Infections, Institut Pasteur, Paris | Marion Barbet, Sylvie Behillil, Frédéric Lemoine, Corinne Maufrais, Christophe Malabat, Méline Bizard, Angela Brisebarre, Camille Capel, Louise Lefrançois, Etienne Simon-Lorière, Vincent Enouf, Maud Vanpeene, Sylvie van der Werf,JéRome Guinard |
| EPI_ISL_1621700 | Hospital | National Reference Center for Viruses of Respiratory Infections, Institut Pasteur, Paris | Marion Barbet, Sylvie Behillil, Frédéric Lemoine, Corinne Maufrais, Christophe Malabat, Méline Bizard, Angela Brisebarre, Camille Capel, Louise Lefrançois, Etienne Simon-Lorière, Vincent Enouf, Maud Vanpeene, Sylvie van der Werf,Nabil Gastli |
| EPI_ISL_1621793 | Hospital | National Reference Center for Viruses of Respiratory Infections, Institut Pasteur, Paris | Marion Barbet, Sylvie Behillil, Frédéric Lemoine, Corinne Maufrais, Christophe Malabat, Méline Bizard, Angela Brisebarre, Camille Capel, Louise Lefrançois, Etienne Simon-Lorière, Vincent Enouf, Maud Vanpeene, Sylvie van der Werf,Jean-Philippe Emond |
| EPI_ISL_1625614 | The Jackson Laboratory | The Jackson Laboratory | Bergeron D, Renzette N, Adams M, Omerza G, Kelly K, Long J, Li L |
| EPI_ISL_1627710 | LABORATORIUM BADA KLINICZNYCH WSSE w OPOLU | 1. National Institute of Public Health - National Institute of Hygiene; 2. Eurofins Genomics Europe Sequencing GmbH | Wokowicz Tomasz, Zacharczuk Katarzyna, Sadkowska-Todys Magorzata, Gierczyki Rafa, Eurofins Genomics Europe Sequencing Team, ECDC COVID-19 WGS support team |
| EPI_ISL_1627967 | LABORATORIA MEDYCZNE OPTIMED | 1. National Institute of Public Health - National Institute of Hygiene; 2. Eurofins Genomics Europe Sequencing GmbH | Wokowicz Tomasz, Zacharczuk Katarzyna, Sadkowska-Todys Magorzata, Gierczyki Rafa, Eurofins Genomics Europe Sequencing Team, ECDC COVID-19 WGS support team |
| EPI_ISL_1629490, EPI_ISL_1629499, EPI_ISL_1629509, EPI_ISL_1629585, EPI_ISL_1629694 | Clinical Microbiology, Infection Prevention and Control | Section for Molecular Diagnostics | Björn Hallström, Jonas Björkman |
| EPI_ISL_1629709 | Infectious Disease Diagnostics laboratory at the Children's Hospital of Philadelphia | Planet Lab, Children's Hospital of Philadelphia | Ahmed M. Moustafa, Colleen Bianco, Lidiya Denu, Azad Ahmed, Rebecca M. Harris, Josh Chang Mell, Paul J. Planet |
| EPI_ISL_761305, EPI_ISL_782148 | Lighthouse Lab in Alderley Park | Wellcome Sanger Institute for the COVID-19 Genomics UK (COG-UK) Consortium | Jacquelyn Wynn, Mairead Hyland, The Lighthouse Lab in Alderley Park and Alex Alderton, Roberto Amato, Sonia Goncalves, Ewan Harrison, David K. Jackson, Ian Johnston, Dominic Kwiatkowski, Cordelia Langford, John Sillitoe on behalf of the Wellcome Sanger Institute COVID-19 Surveillance Team |
| EPI_ISL_822694 | Wales Specialist Virology Centre Sequencing lab: Pathogen Genomics Unit | COVID-19 Genomics UK (COG-UK) Consortium | Catherine Moore, Johnathan Evans, Laura Gifford, Malorie Perry, Simon Cottrell, Angela Marchbank, Alec Birchley, Alexander Adams, Amy Gaskin, Bree Gatica-Wilcox, Jason Coombes, Joel Southgate, Lauren Gilbert, Lee Graham, Nicole Pacchiarini, Sara Kumziene-Summerhayes, Sarah Taylor, Sophie Jones, Sara Rey, Matthew Bull, Joanne Watkins, Sally Corden, Tom Connor |
| EPI_ISL_835003, EPI_ISL_835007, EPI_ISL_835036, EPI_ISL_846335 | Lighthouse Lab in Alderley Park | Wellcome Sanger Institute for the COVID-19 Genomics UK (COG-UK) Consortium | Jacquelyn Wynn, Mairead Hyland, The Lighthouse Lab in Alderley Park and Alex Alderton, Roberto Amato, Sonia Goncalves, Ewan Harrison, David K. Jackson, Ian Johnston, Dominic Kwiatkowski, Cordelia Langford, John Sillitoe on behalf of the Wellcome Sanger Institute COVID-19 Surveillance Team |
| EPI_ISL_851504, EPI_ISL_851507 | Lighthouse Lab in Glasgow | Wellcome Sanger Institute for the COVID-19 Genomics UK (COG-UK) Consortium | Harper VanSteenhouse, Yumi Kasai, David Gray, Carol Clugston, Anna Dominiczak and Alex Alderton, Roberto Amato, Sonia Goncalves, Ewan Harrison, David K. Jackson, Ian Johnston, Dominic Kwiatkowski, Cordelia Langford, John Sillitoe on behalf of the Wellcome Sanger Institute COVID-19 Surveillance Team |
| EPI_ISL_864004, EPI_ISL_879765, EPI_ISL_880971, EPI_ISL_881226, EPI_ISL_881370, EPI_ISL_881376, EPI_ISL_885376 | Lighthouse Lab in Alderley Park | Wellcome Sanger Institute for the COVID-19 Genomics UK (COG-UK) Consortium | Jacquelyn Wynn, Mairead Hyland, The Lighthouse Lab in Alderley Park and Alex Alderton, Roberto Amato, Sonia Goncalves, Ewan Harrison, David K. Jackson, Ian Johnston, Dominic Kwiatkowski, Cordelia Langford, John Sillitoe on behalf of the Wellcome Sanger Institute COVID-19 Surveillance Team |
| EPI_ISL_908202, EPI_ISL_908209 | Lighthouse Lab in Cambridge | Wellcome Sanger Institute for the COVID-19 Genomics UK (COG-UK) Consortium | Rob Howes, The Lighthouse Lab in Cambridge and Alex Alderton, Roberto Amato, Sonia Goncalves, Ewan Harrison, David K. Jackson, Ian Johnston, Dominic Kwiatkowski, Cordelia Langford, John Sillitoe on behalf of the Wellcome Sanger Institute COVID-19 Surveillance Team |
| EPI_ISL_908354, EPI_ISL_908386, EPI_ISL_908474 | Lighthouse Lab in Alderley Park | Wellcome Sanger Institute for the COVID-19 Genomics UK (COG-UK) Consortium | Jacquelyn Wynn, Mairead Hyland, The Lighthouse Lab in Alderley Park and Alex Alderton, Roberto Amato, Sonia Goncalves, Ewan Harrison, David K. Jackson, Ian Johnston, Dominic Kwiatkowski, Cordelia Langford, John Sillitoe on behalf of the Wellcome Sanger Institute COVID-19 Surveillance Team |
| EPI_ISL_920689, EPI_ISL_920690 | University College London Hospital | COVID-19 Genomics UK (COG-UK) Consortium | Judith Heaney, Matthew Byott, Catherine Houlihan, Dan Frampton, Stuart Kirk, Moira Spyer and Eleni Nastouli |
| EPI_ISL_932020, EPI_ISL_932032 | Lighthouse Lab in Alderley Park | Wellcome Sanger Institute for the COVID-19 Genomics UK (COG-UK) Consortium | Jacquelyn Wynn, Mairead Hyland, The Lighthouse Lab in Alderley Park and Alex Alderton, Roberto Amato, Sonia Goncalves, Ewan Harrison, David K. Jackson, Ian Johnston, Dominic Kwiatkowski, Cordelia Langford, John Sillitoe on behalf of the Wellcome Sanger Institute COVID-19 Surveillance Team |
| EPI_ISL_938238 | Lighthouse Lab in Cambridge | Wellcome Sanger Institute for the COVID-19 Genomics UK (COG-UK) Consortium | Rob Howes, The Lighthouse Lab in Cambridge and Alex Alderton, Roberto Amato, Sonia Goncalves, Ewan Harrison, David K. Jackson, Ian Johnston, Dominic Kwiatkowski, Cordelia Langford, John Sillitoe on behalf of the Wellcome Sanger Institute COVID-19 Surveillance Team |
| EPI_ISL_939292 | Lighthouse Lab in Milton Keynes | Wellcome Sanger Institute for the COVID-19 Genomics UK (COG-UK) Consortium | The Lighthouse Lab in Milton Keynes and Alex Alderton, Roberto Amato, Sonia Goncalves, Ewan Harrison, David K. Jackson, Ian Johnston, Dominic Kwiatkowski, Cordelia Langford, John Sillitoe on behalf of the Wellcome Sanger Institute COVID-19 Surveillance Team |
| EPI_ISL_945381 | Lighthouse Lab in Alderley Park | Wellcome Sanger Institute for the COVID-19 Genomics UK (COG-UK) Consortium | Jacquelyn Wynn, Mairead Hyland, The Lighthouse Lab in Alderley Park and Alex Alderton, Roberto Amato, Sonia Goncalves, Ewan Harrison, David K. Jackson, Ian Johnston, Dominic Kwiatkowski, Cordelia Langford, John Sillitoe on behalf of the Wellcome Sanger Institute COVID-19 Surveillance Team |
| EPI_ISL_957114 | Lighthouse Lab in Glasgow | Wellcome Sanger Institute for the COVID-19 Genomics UK (COG-UK) Consortium | Harper VanSteenhouse, Yumi Kasai, David Gray, Carol Clugston, Anna Dominiczak and Alex Alderton, Roberto Amato, Sonia Goncalves, Ewan Harrison, David K. Jackson, Ian Johnston, Dominic Kwiatkowski, Cordelia Langford, John Sillitoe on behalf of the Wellcome Sanger Institute COVID-19 Surveillance Team |
| EPI_ISL_958243 | Lighthouse Lab in Milton Keynes | Wellcome Sanger Institute for the COVID-19 Genomics UK (COG-UK) Consortium | The Lighthouse Lab in Milton Keynes and Alex Alderton, Roberto Amato, Sonia Goncalves, Ewan Harrison, David K. Jackson, Ian Johnston, Dominic Kwiatkowski, Cordelia Langford, John Sillitoe on behalf of the Wellcome Sanger Institute COVID-19 Surveillance Team |
| EPI_ISL_969661 | Lighthouse Lab in Alderley Park | Wellcome Sanger Institute for the COVID-19 Genomics UK (COG-UK) Consortium | Jacquelyn Wynn, Mairead Hyland, The Lighthouse Lab in Alderley Park and Alex Alderton, Roberto Amato, Sonia Goncalves, Ewan Harrison, David K. Jackson, Ian Johnston, Dominic Kwiatkowski, Cordelia Langford, John Sillitoe on behalf of the Wellcome Sanger Institute COVID-19 Surveillance Team |
| EPI_ISL_980572 | Lighthouse Lab in Milton Keynes | Wellcome Sanger Institute for the COVID-19 Genomics UK (COG-UK) Consortium | The Lighthouse Lab in Milton Keynes and Alex Alderton, Roberto Amato, Sonia Goncalves, Ewan Harrison, David K. Jackson, Ian Johnston, Dominic Kwiatkowski, Cordelia Langford, John Sillitoe on behalf of the Wellcome Sanger Institute COVID-19 Surveillance Team |
| EPI_ISL_987108, EPI_ISL_988812 | Lighthouse Lab in Alderley Park | Wellcome Sanger Institute for the COVID-19 Genomics UK (COG-UK) Consortium | Jacquelyn Wynn, Mairead Hyland, The Lighthouse Lab in Alderley Park and Alex Alderton, Roberto Amato, Sonia Goncalves, Ewan Harrison, David K. Jackson, Ian Johnston, Dominic Kwiatkowski, Cordelia Langford, John Sillitoe on behalf of the Wellcome Sanger Institute COVID-19 Surveillance Team |

| | | | |
|---|---|---|---|
| | | | (http://www.sanger.ac.uk/covid-team) |
| EPI_ISL_991123, EPI_ISL_991635, EPI_ISL_994481 | Lighthouse Lab in Alderley Park | Wellcome Sanger Institute for the COVID-19 Genomics UK (COG-UK) Consortium | Jacquelyn Wynn, Mairead Hyland, The Lighthouse Lab in Alderley Park and Alex Alderton, Roberto Amato, Sonia Goncalves, Ewan Harrison, David K. Jackson, Ian Johnston, Dominic Kwiatkowski, Cordelia Langford, John Sillitoe on behalf of the Wellcome Sanger Institute COVID-19 Surveillance Team |
| EPI_ISL_995070 | Pandemic Response Lab - NYC | Pandemic Response Lab, R&D | Henry Lee, Michael Hammerling, Melissa Hopkins, Cybill del Castillo, William Ward, Pradeep Bugga, Haiping Hao, Jon Laurent |
| EPI_ISL_996136, EPI_ISL_996160 | Quadram Institute Bioscience | COVID-19 Genomics UK (COG-UK) Consortium | Dave J. Baker, Gemma L. Kay, Alp Aydin, Thanh Le-Viet, Steven Rudder, Ana P. Tedim, Anastasia Kolyva, Maria Diaz, Leonardo de Oliveira Martins, Nabil-Fareed Alikhan, Lizzie Meadows, Rachael Stanley, Ngozi Elumogo, Muhammed Yasir, Nicholas M. Thomson, Alexander J Trotter, Rachel Gilroy, Samuel Bloomfield, Claire Stuart, Andrew Bell, Reenesh Prakash, Samir Dervisevic, Alison E. Mather, John Wain, Mark Webber, Andrew J. Page, Justin O'Grady |
| EPI_ISL_996993, EPI_ISL_996995 | Department of Pathology, University of Cambridge | COVID-19 Genomics UK (COG-UK) Consortium | Aminu S. Jahun, Yasmin Chaudhry, Iliana Georgana, Myra Hosmillo, Rhys Izuagbe, William L. Hamilton, Martin D. Curran, Surendra Parmar, Ian Goodfellow |
